# Upper Limit for Angular Compounding Speckle Reduction

**DOI:** 10.1101/239350

**Authors:** Yonatan Winetraub, Chris Wu, Steven Chu, Adam de la Zerda

**Affiliations:** Biophysics Program at Stanford, Stanford, California 94305, USA; Department of Structural Biology, Stanford University, Stanford, California 94305, USA; Molecular Imaging Program at Stanford, Stanford, California 94305, USA; The Bio-X Program, Stanford, California 94305, USA; Departments of Physics and Molecular and Cellular Physiology, Stanford University, Stanford, California 94305, USA; The Chan Zuckerberg Biohub, San Francisco, California 94158, USA

**Keywords:** Optical Coherence Tomography, Speckle Reduction, Angular Compounding

## Abstract

Previous studies of angular compounding for speckle reduction in optical coherence tomography may not have fully accounted for optical aberrations, which produce unintended spatial averaging and concomitant loss of spatial resolution. We accounted for such aberrations by aligning our system and measuring distortions in the images, and found that speckle reduction by angular compounding was limited. Our theoretical analysis using Monte Carlo simulations indicates that “pure” angular compounding over 13° (our full numerical aperture) can improve the signal-to-noise ratio by no more than a factor of 1.5, significantly lower than values reported in literature. Analysis suggests that illuminating only part of the lens to further reduce speckle also involves a trade-off with resolution roughly equivalent to spatial averaging. We conclude that angular compounding provides substantially less benefit than previously expected.

---

Optical coherence tomography (OCT) is a powerful tool for non-invasive probing of the microstructure of biological tissue. Because the technique relies on coherent detection of scattered light, however, OCT images are confounded by speckle noise: a grainy texture that reduces the signal-to-noise ratio (SNR) and the effective spatial resolution. A widely used method to reduce speckle noise is angular compounding [12], which averages results obtained with the imaging beam probing the sample at different angles. Both the imaging system and sample can remain stationary during the scan, allowing high imaging throughput with high image quality [9, 5]. Angular compounding has been reported to significantly decrease speckle, with the SNR of angular compounded images as much as 6.5 times that of a corresponding single-angle image [5]. Furthermore, several studies [1, 4] found angular compounding to increase SNR significantly more than spatial averaging. Recent work has suggested combining image processing with angular compounding [3, 6] to further reduce speckle. A key benefit claimed for angular compounding (e.g., compared to spatial averaging) is that it may achieve speckle reduction with minimal to no degradation of spatial resolution [12, 9, 5, 1, 4]. As angular compounding becomes more widely used, it is important to study how it differs from spatial averaging, and to what extent it removes speckle without impairing spatial resolution.

In this work, we employed a common setup [Fig. 1(a)] for angular compounded OCT, in which a galvo-controlled scanning mirror is offset a varying distance *h* from the optical axis [4]. In classical OCT, the galvo mirror remains centered on the optical axis (i.e., *h* = 0). To acquire a single pixel of an A-scan, a ray reflects off the galvo mirror, passes through the optical system at point (*s, θ*), and scatters from the sample at point (*x, y, z*), as depicted in Fig. 1(b). The full A-scan is built up from a series of such pixels due to the ray scattering at different depths z. To acquire a B-scan, the galvo mirror rotates, sweeping s from −1 to 1 while keeping *θ* constant. Varying the distance *h* provides scans of (*x, y, z*) using rays at different angles, which are averaged for the angular compounded image [4].

**Figure 1:**
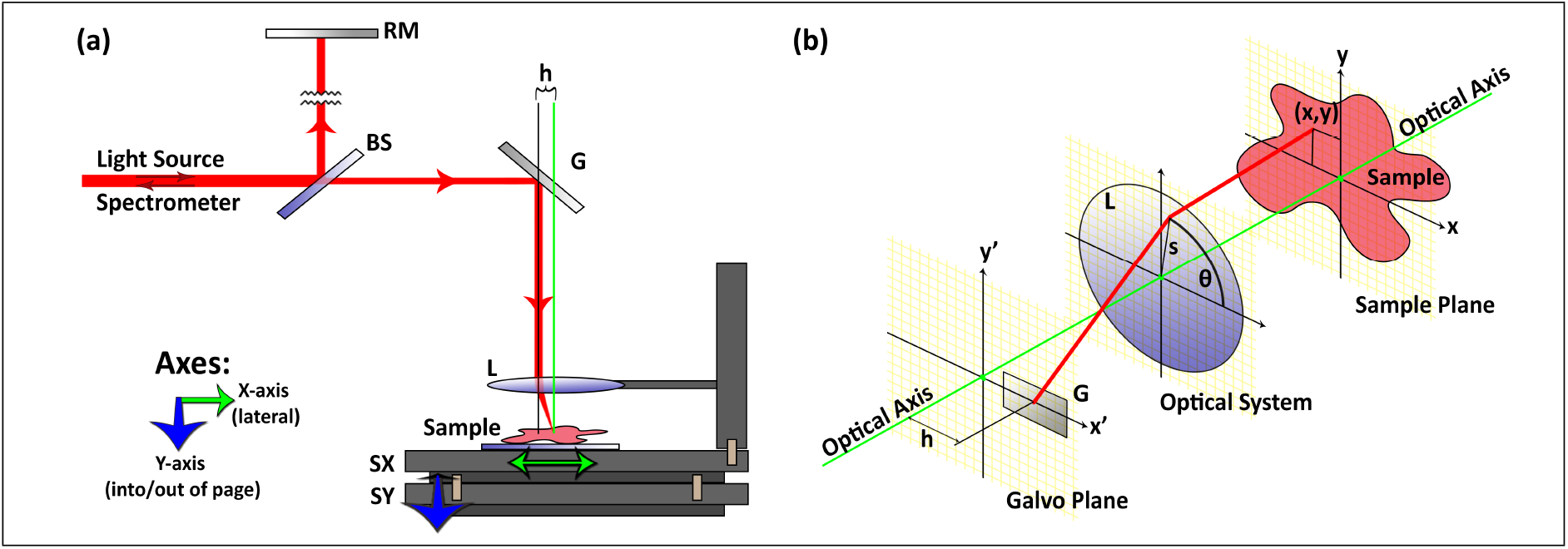
(a) Angle-resolved OCT system schematic. RM, reference mirror; BS, beam splitter; G, galvo-controlled scanning mirror; L, objective lens (*f* = 18 mm); SX, SY, translation stages moving sample and optics together with respect to laser source. (b) Sample arm optics showing ray trajectory from galvo mirror to sample. A ray reflects off the galvo mirror at point (*x′* = *h*, *y′* = 0), passes through the optical system’s aperture at point (*s*, *θ*), and is scattered by the sample at point (*x, y, z*).

Using a lens model with axially symmetrical aberrations, we can calculate (*x, y*) as a function of *h*, *s*, *θ* [13]:

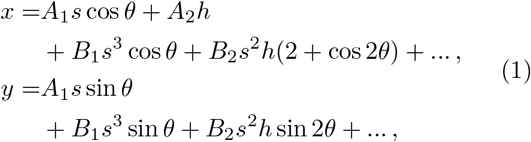

where constants *A_1_*, *A_2_* describe the first-order imagery; and *B_1_*, *B_2_* describe primary aberrations. Both *x* and *y* depend on *h*. Thus, changes in *h* not only result in reflection of light from the sample at different angles (angular compounding) but also move the beam with respect to the sample, which introduces spatial averaging. A “pure” angular compounding setup should correct for these displacements.

Recent angular compounding work has suggested the use of image registration by global translation estimation prior to averaging, to reduce spatial averaging [10, 2, 3]. These corrections do remove some of the spatial averaging, but are insufficient since *x* and *y* are nonlinear functions of *h* and *s*. Furthermore, *h* introduces distortions out of the B-scan plane, so a full 3D volume scan is required to perform image registration.

To account for aberrations in our model, we first align the galvo B-scan direction with the *x′*-axis (i.e., set *θ* = 0). This confines distortions to the B-scan plane (now the *x-z* plane), simplifying Eq. 1:

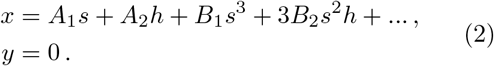

We then introduce non-axially symmetrical aberrations to the model, and neglect higher order terms, writing the result in terms of the system’s “distortion field,” *U*, *W*, *V*:

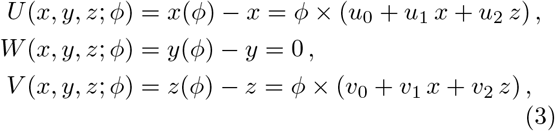

where *φ* = tan^−1^(*h/f*); *f* is the objective focal length; and the *u_i_*, *v_i_* are aberration parameters summarizing all optical distortions, which are unique to each optical setup. We also use the notation *x* = *x*(*φ* = 0), *y* = *y*(*φ* = 0), *z* = *z*(*φ* = 0). To understand the true extent of speckle reduction due to pure angular compounding, we must account for this distortion field.

To test our model experimentally, we proceeded as follows: First, we used a V target to align the B-scan direction to *θ* = 0. Next, we imaged a phantom sample with standard angular compounding, and measured the distortion field. Finally, we used the distortion field to obtain a corrected angular compounded image of the phantom. In all experiments, we used a commercial spectral domain OCT system (Ganymede HR SD-OCT using LSM02-BB lens, ThorLabs, Newton, NJ) with 4 *μm* FWHM optical pixel size. The SD-OCT light source was a superluminescent diode (SLD) with center wavelength *λ* = 900 nm and spectral bandwidth 200 nm. As a preprocessing step, we used averages of 20 B-scans to remove photon shot noise from the data.

In the alignment step, our goal was to align *θ* = 0 to an accuracy of one optical pixel over the course of a 500 μm scan (i.e., Δ*θ* = 0.46°). We fabricated a V target on a silicon wafer using standard lithographic and dry etching processes. The target consisted of two perpendicular trenches, each 50 μm wide × 28 μm deep [Fig. 2(a)]. We mechanically fixed the V target to a translation stage along the *y*-axis. We applied a few microliters of gold nanorod solution (OD 50) [14] to increase contrast.

**Figure 2:**
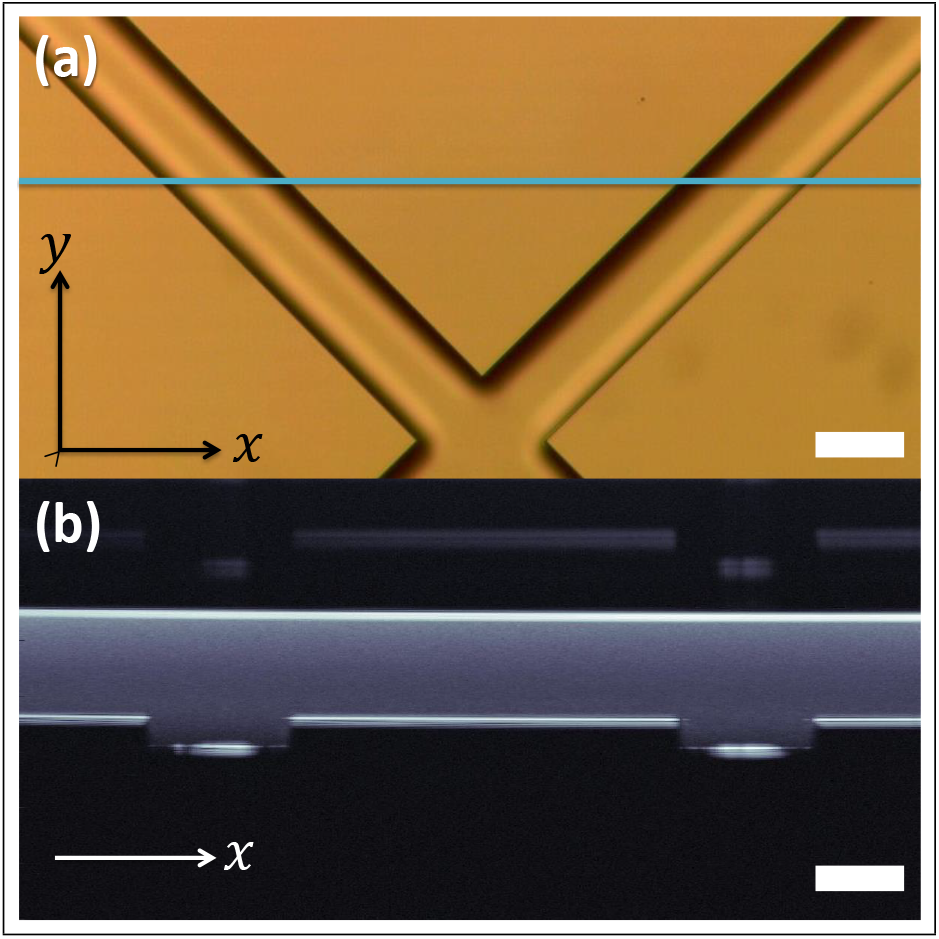
(a) Bright field microscope image of calibration target showing two trenches forming a V. (b) OCT B-scan taken along the blue line in (a). Alignment of the B-scan with the *x*-axis is measured by observing the movement of the trench centers when the target is moved along the *y*-axis. Scale bars: 50 μm.

We imaged the V target such that both trenches were visible in the B-scan [Fig. 2(b)]. We estimated the center position of each trench and observed how these changed when the V target was moved along the *y*-axis, and adjusted our system accordingly. When the centers moved by equal and opposite amounts during *y*-translation, the system was aligned to *θ* = 0.

After alignment, we imaged a 2% w/v Intralipid phantom (made from 20% w/v stock solution in agarose) at 17 angles φ,

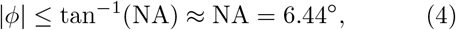

where NA is the numerical aperture of our system. The angles were at equal intervals of 0.8°. Fig. 3(a) shows an OCT B-scan of the phantom acquired from a single angle. The speckle contrast is defined as *σ_I_*/〈*I*〉, the standard deviation of the intensity over the mean linear intensity, in an area of uniform scattering. The SNR is the inverse of this quantity, 〈*I*〉/*σ_I_*. When we performed standard angular compounding using all 17 angles [Fig. 3(b)], the SNR increased by a factor of 2.70 compared to a singleangle image. (We call this a relative SNR, or RSNR, of 2.7.) This result is comparable to values reported elsewhere.

**Figure 3:**
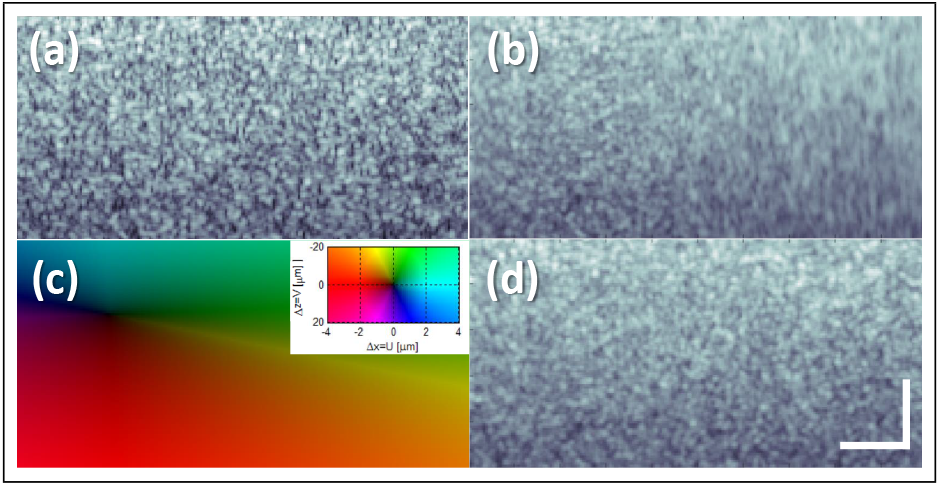
Angular compounding of Intralipid phantom. (a) B-scan image (*x-z* plane) acquired from a single angle. Relative SNR (RSNR) = 1, by definition. (b) Image obtained by standard angular compounding of scans at 17 angles *φ*, with |*φ*| ≤ 6.4°. RSNR = 2.7. (c) Estimated distortion field at *φ* = 6.4° shows significant movement in both *x* and *z* directions. Inset depicts 2D color scale for the displacements. (d) Image using angular compounding corrected for the distortion field. RSNR is significantly reduced to 1.65. Scale bars: 30 *μ*m.

We estimated our system’s distortion field by measuring the translational displacement of 100 random patches (each 30 *μ*m × 30 *μ*m) in our images using a subpixel registration algorithm [7], then fitting the patch displacements to Eq. 3 using least squares to estimate *u*_0_, *u*_1_, *u*_2_, *v*_0_, *v*_1_, *v*_2_. As can be seen in Fig. 3(c), the distortion field was significant (as large as 12 *μ*m) and very different from a translation-only registration error. After this distortion was accounted for [Fig. 3(d)], the RSNR was reduced to 1.65. The residual error of the patch motion fit was ~1-2 *μ*m, suggesting that small uncorrected displacements remain in the data.

We noticed that in the dark area in Fig. 3(c), near the lens’s optical axis, *U* and *V* have values less than 500 nm. In an attempt to obtain a corrected image with even less residual distortion than that of Fig. 3(d), we used this region to acquire 17 B-scans along the *y-z* plane (*θ* = 90°). The RSNR, comparing Figs. 4(a) and 4(b), was further reduced to 1.49. On examining small regions across a series of single-angle images [e.g., Fig. 4(c)], we saw that the speckle pattern did not change significantly when *φ* changed by relatively large angles. These results suggest that even the slightest unwanted translation can reduce speckle, and studies of pure angular compounding ought to account for movements of ~1/10 of a pixel (much lower than the limit suggested by [8]). Conversely, by deliberately using small subpixel shifts, spatial compounding alone might significantly reduce speckle with only slight loss of resolution.

**Figure 4:**
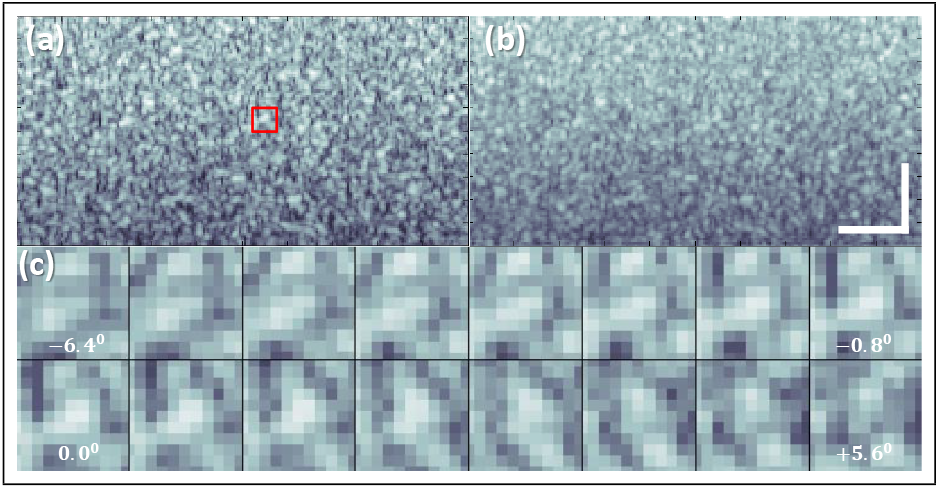
Angular compounding of Intralipid phantom at position where the distortion field is <500 nm. (a) B-scan image (*y-z* plane) acquired from a single angle. (b) Image obtained by standard angular compounding of 17 scans. RSNR = 1.49, very close to our theoretical limit. (c) Area bounded by red box in (a), enlarged for angles −6.4° to −0.8° (top row) and 0° to 5.6° (bottom row). The similarity of the pattern at many angles explains why compounding achieves only modest speckle reduction. Scale bars: 30 *μ*m.

We now model speckle contrast assuming that the distortion field is accounted for, to derive a theoretical upper limit for speckle reduction by pure angular compounding for our system. We have previously shown [15] that a simple model assuming *N* identical isotropic scatterers randomly distributed in an imaging voxel of size (*σ_x_*, *σ_y_*, *σ_z_*) can describe speckle behavior in OCT images. We use coordinates (*x*, *y*, *z*) centered on the voxel’s center. When a point scatterer at (*x, y, z*) scatters light, it generates an electric field with real *g* and imaginary *h* components. Here we analyze *g*; the analysis for *h* is similar.

First, note that

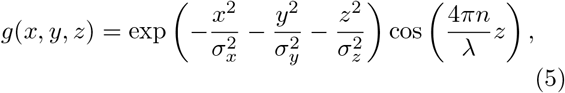

where *λ* is the OCT central wavelength, and n is index of refraction. When the imaging beam rotates by an angle *φ*, a scatterer’s position in the voxel-based coordinates (after accounting for global displacements) will change as follows:

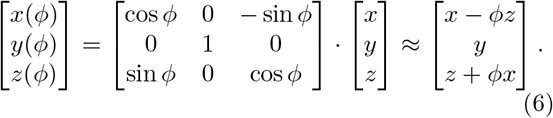

We use Eqs. 5 and 6 to generate a Monte Carlo simulation (randomizing the scatterer position) to estimate Cov[*g*(*φ*),*g*(*φ* = 0)]. Surprisingly, the correlation *R* as a function of *φ* computed with this simulation fits very well with a Gaussian form:

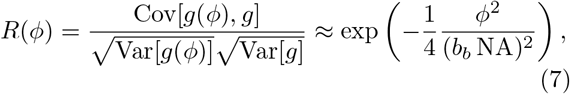

where *b_b_* = 0.372 from the simulation. Furthermore, *R(φ)* drops to 0 only at very high angles. An intuitive explanation is that in order to significantly change *g*, the scatterer must move a considerable distance, to change its intensity and phase. For small angles, however, the position (*x, z*) changes only as (*φz*, *φx*), which is too small compared to voxel size.

Next, we perform a simulation with *N* independent scatterers in the voxel, summing the electric fields that they generate to compute the voxel’s real electric field *G*:

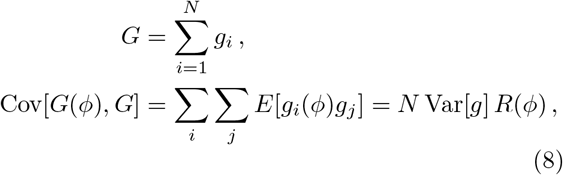

and similarly for the imaginary field *H*. In other words, we find that the correlation between *G(φ)* and *G* is the same as that of individual particles: *R(φ)*.

Finally, we compute the covariance of the speckle linear intensity, Cov[*I(φ), I*], assuming *G*, *H* are independent Gaussian variables, so that 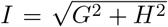 follows a Rayleigh distribution. This in turn yields the speckle correlation, *ρ(φ)*:

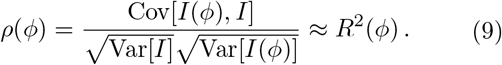

Substituting Eq. 7 into Eq. 9, we get

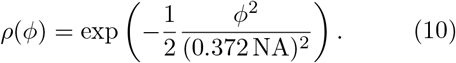

Next, we estimate the angular compounding signal *B*. In an ideal case, we could average the signal acquired from all possible angles, attenuated by the optical support of the system:

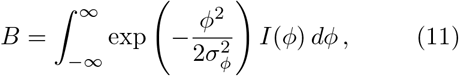

where *σ_φ_* = 0.5 NA, as NA is the 1/*e*^2^ radius of the lens. Finally, we compute the relative SNR for this ideal case, which we call 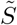:

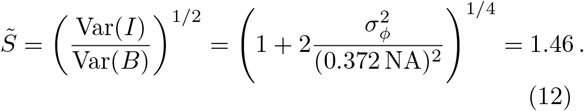

We conclude that 1.46 is the upper limit for the RSNR due to pure angular compounding. Any additional speckle reduction observed should be attributed to spatial averaging. Note that 1.46 is close to 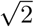, the RSNR to be expected by averaging just 2 completely uncorrelated speckle patterns.

Up to this point, we have assumed that the collimated laser beam in Fig. 1(a) illuminates the full width of the lens. Speckle reduction can be further increased, however, by illuminating a small part of the lens, which increases the ratio of the compounding angles to the effective NA of the illuminated area. Assuming the relative illuminated area is *p*^2^, then *b_b_* in Eq. 7 changes to *b_b_p*, leading to an RSNR limit of:

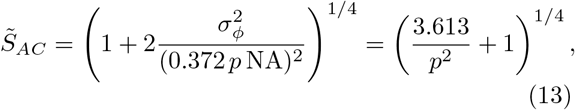

which can be higher than 1.46. However, since the partial illumination of the lens also reduces the spatial resolution by a factor of *p*, we should compare the angular compounding scenario to an alternative in which we fully illuminate the lens, but perform spatial averaging that reduces spatial resolution by the same factor *p*. We can repeat Eqs. 7 to 10 using a displacement-based model and Monte Carlo simulation, or use similar analysis [11], to conclude:

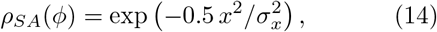

where *x*/*σ_x_* is the amount of spatial averaging compared to pixel size. Similar to Eq. 11, we assume a Gaussian smoothing of width 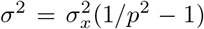 to match resolution loss between angular compounding and spatial averaging. As a result, Eq. 13 is transformed to

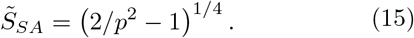

Comparing Eqs. 13 and 15, we find that 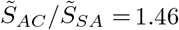 at *p* = 1 and declines as *p* is reduced. Note that all our analysis above was performed for single-axis compounding.

In conclusion, angular compounding adds only slight value compared to simple post-processing spatial averaging. Greater speckle reduction seemingly by angular compounding may be the result of unintended spatial averaging due to lens aberrations. We have described two steps to measure and account for such aberrations: scan alignment using a V target, and distortion field estimation. Once distortion is accounted for, the pure angular compounding that remains (for single-axis scanning over 13°, the full NA) is expected to increase SNR by no more than a factor of about 1.5. We hope that future angular compounding research will use our method to ensure that unintended displacements are smaller than ~1/10 of a pixel. Such correction will yield higher spatial resolution and reliable speckle reduction figures. We also recommend reporting the illumination fraction of the lens. Partial illumination may increase speckle reduction, but only at the expense of reduced spatial resolution, comparable to the cost of equivalent spatial averaging at full illumination.

## Acknowledgments

A.d.l.Z is a Chan Zuckerberg Biohub investigator and a Pew-Stewart Scholar for Cancer Research supported by The Pew Charitable Trusts and The Alexander and Margaret Stewart Trust. Y.W. is grateful for a Stanford Bowes Bio-X Graduate Fellowship, Stanford Biophysics Program training grant (T32 GM-08294). Authors would like to acknowledge Stanford Nanofabrication Facility (SNF) faculty and staff for providing help and guidance in design, manufacturing, and testing of V target; Edwin Yuan and Elliott SoRelle for comments and discussion; and Graham P. Collins for manuscript editing.

